# Methods for evaluating unsupervised vector representations of genomic regions

**DOI:** 10.1101/2023.08.28.555137

**Authors:** Guangtao Zheng, Julia Rymuza, Erfaneh Gharavi, Nathan J. LeRoy, Aidong Zhang, Nathan C. Sheffield

## Abstract

Representation learning models have become a mainstay of modern genomics. These models are trained to yield vector representations, or embeddings, of various biological entities, such as cells, genes, individuals, or genomic regions. Recent applications of unsupervised embedding approaches have been shown to learn relationships among genomic regions that define functional elements in a genome. Unsupervised representation learning of genomic regions is free of the supervision from curated metadata and can condense rich biological knowledge from publicly available data to region embeddings. However, there exists no method for evaluating the quality of these embeddings in the absence of metadata, making it difficult to assess the reliability of analyses based on the embeddings, and to tune model training to yield optimal results. To bridge this gap, we propose four evaluation metrics: the cluster tendency score (CTS), the reconstruction score (RCS), the genome distance scaling score (GDSS), and the neighborhood preserving score (NPS). The CTS and RCS statistically quantify how well region embeddings can be clustered and how well the embeddings preserve information in training data. The GDSS and NPS exploit the biological tendency of regions close in genomic space to have similar biological functions; they measure how much such information is captured by individual region embeddings in a set. We demonstrate the utility of these statistical and biological scores for evaluating unsupervised genomic region embeddings and provide guidelines for learning reliable embeddings.

**Availability:** Code is available at https://github.com/databio/geniml

## Introduction

Genomic regions, or intervals, define functional elements in the genome, such as enhancers, promoters, or transcription factor binding sites (1). A region is represented as a pair of coordinates marking its location on the genome. A set of such regions, often stored in Browser Extensible Data (BED) format, can be used to represent an epigenomics experiment that identifies locations of interest produced by a biological experiment, such as ChIP-Seq (2) or ATAC-Seq (3, 4). Through broad effort (1, 5), the amount of epigenome data has rapidly increased, and almost 100,000 BED files are now available on the NCBI Gene Expression Omnibus (GEO) (6–8). This growing resource of BED files contains rich biological information and have been used to interpret the human genome and drive discovery in genetic variation and gene regulation (9–12). However, the ever-increasing volume of genomic region data also makes region-based analyses computationally costly, since analysis often requires computing region overlaps among region sets (13–15).

We recently introduced region-set2vec (16), an unsupervised method to learn representations for genomic region sets. The method learns low-dimensional vectors called embeddings as representations for genomic region sets, using only the region co-occurrence information contained in region sets. With region-set2vec, we can leverage the publicly available genomic region data, even without annotations, and can replace some costly region overlap calculation with more efficient vector similarity measures. To create region set embeddings, region-set2vec first must learn region embeddings, which are then averaged to represent the region set. We previously demonstrated the value of region set embeddings. We reason that the underlying region embeddings could be useful independently; for example, we can infer the function of an unknown region based on its closeness to other known regions in the embedding space, or use them for fast annotation and clustering on single-cell data (17). To develop such methods and concepts further, we first sought a way to evaluate region embeddings independently and objectively. Our previous study evaluated region set embeddings, but no method has yet been proposed to evaluate the quality of the underlying region embeddings.

To bridge this gap, we propose four novel statistical and biological scores to measure the quality of region embeddings. First, we propose two statistical scores that use only the training data and the region embeddings: 1) the cluster tendency score (CTS), which measures how well a set of region embeddings can form clusters; and 2) the reconstruction score (RCS), which tests whether a region embedding can reconstruct the training data. Next, we propose two biologically-motivated scores that exploit the tendency of regions near one another on the genome to have similar biological functions. We reasoned that embedding distances that reflect functional similarity between regions should be biased toward smaller embedding distances for regions nearby on the genome. In other words, two embeddings close in latent space should be, on average, closer in genome space. To assess whether region embeddings capture this tendency, we propose the genome distance scaling score (GDSS) and the neighborhood preserving score (NPS). The GDSS shows the degree to which the embedding distance between two regions scales with genome distance. We expect to obtain a positive and large scaling value if the embeddings recapitulate genome distance. The NPS measures how much the neighborhood of a region in the genome space overlaps with the neighborhood of the same region in the embedding space.

These four scores are generally applicable to any set of region embeddings, including Region2Vec embeddings. To demonstrate, we calculated the four scores for three different types of embeddings: 1) Binary embeddings; 2) dimensionality-reduced versions of Binary embeddings; and 3) Region2Vec embeddings trained with various learning parameters. Our results show the advantage of Region2Vec embeddings in capturing the desired biological knowledge with low dimensions, providing new evidence for the value of region-set2vec embeddings (16). Using a classification task, we showed how these scores can reflect the utility of embeddings on a downstream task. Finally, our study provides guidelines for training hyperparameters for learning reliable genomic region embeddings based on the obtained scores.

## Methods

### Data overview

To guide our embedding evaluation methods development, we first collected a representative collection of region sets consisting of 690 transcription factor binding BED files from the ENCODE Uniform TFBS composite track as a region set collection to generate and evaluate region embeddings.

### Tokenization of BED files into universe regions

To compare diverse region sets, many epigenome analysis methods, including our embedding methods, require first re-defining the raw regions into consensus regions. We refer to the consensus region set as a “universe”, and the process of re-defining the raw regions as “tokenization” (Figure 1A). Tokenization allows us to use a shared vocabulary of genomic regions. We tokenize by representing each original region as any universe regions that it overlaps. Original regions that do not overlap any universe region are discarded. Thus, after tokenization, tokenized BED files contain only unique universe regions. As a result, we need only consider learning embeddings for regions in the universe instead of original regions. Tokenization is the first step for any region embedding learning methods considered in this paper.

**Figure 1.**
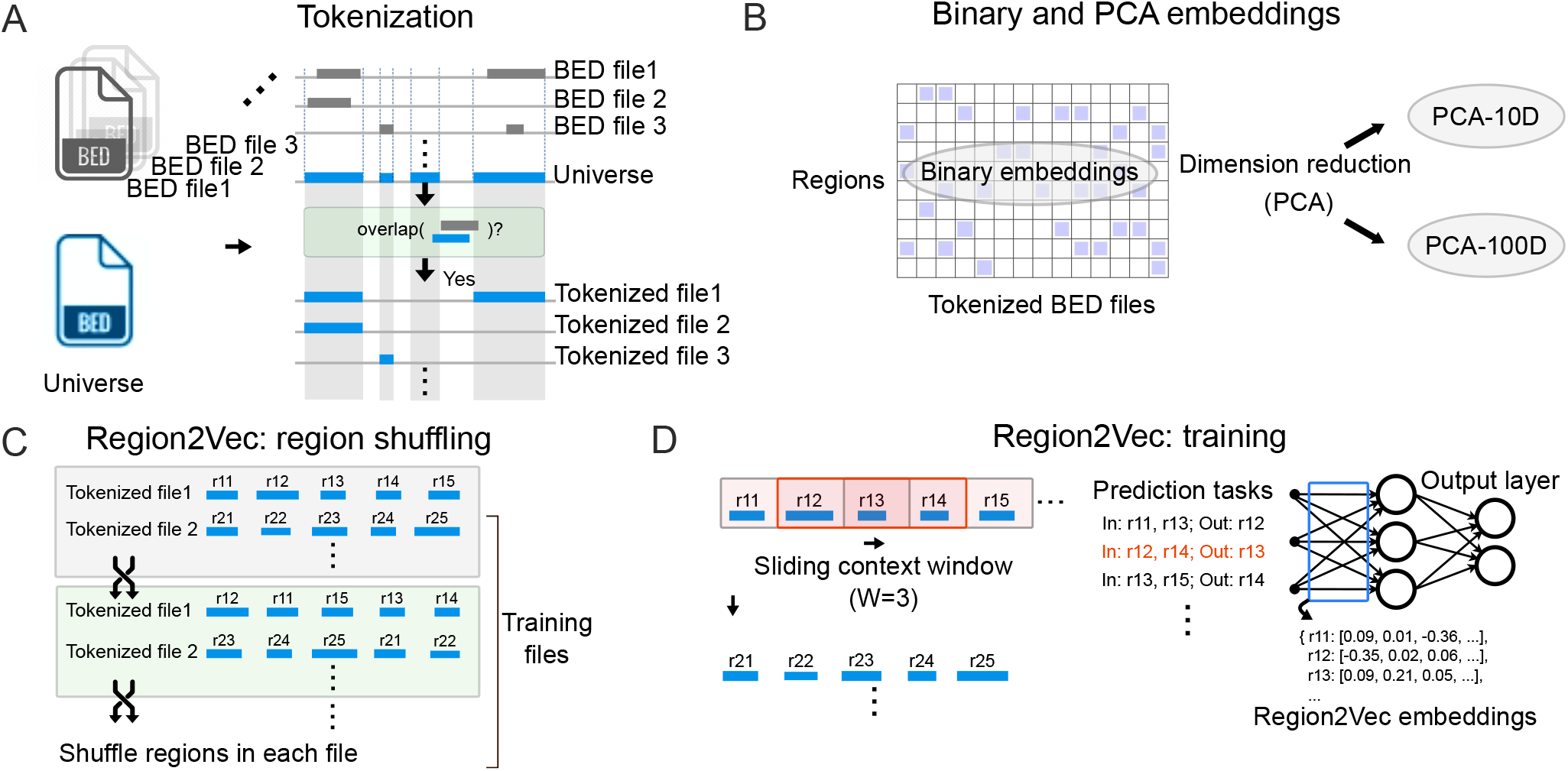
Illustration of unsupervised region embedding generation. **A**. Replace regions in BED files with overlapped regions in a universe. This is called tokenization. **B**. Binary embeddings and the embeddings after keeping 10 (PAC-10D) and 100 (PCA-100D) principal components of the Binary embeddings. **C**. Region2Vec shuffles regions in tokenized BED files to generate more training files. **D**. Region2Vec trains the model using the prediction tasks created by a sliding context window with length W.

To test how our proposed evaluation metrics are affected by region size, number of elements, and fit of the universe, we built seven universes with different numbers of regions and tokenized the 690 BED files into each (Table S2). These include 3 Merge universes, which we built by merging all regions in the collection using different distance thresholds. For example, For the Merge (100) universe, we merged all regions that were less than 100 bp apart. The other 4 universes are not derived from the BED file collection; the Tiling universes are created by simple tiling of fixed-size windows across the genome, and the DHS universe is an external universe defined as regulatory regions from DNase I hypersensitive sites (DHSs) with similar tissue specificity of DNase-seq signal patterns (18). For these non-data-driven universes, we filtered out any regions that do not appear in the tokenized BED file collection. For each universe, we generated region embeddings by first tokenizing the region sets. We then used the tokenized region sets as training data to produce embeddings.

### Constructing region embeddings

We produced region embeddings using three unsupervised methods: 1) Binary embeddings, which are 690-dimensional vectors where each dimension represents presence of a region in one of the 690 BED files; 2) principal component analysis (PCA) embeddings, constructed as the top 10 (PCA-10D) or 100 (PCA-100D) principal components of the binary embedding matrix(19); and 3) Region2Vec embeddings, using 12 different parameter sets for training the Region2Vec model.

#### Binary and PCA embeddings

The simplest embeddings, binary region embeddings, are parameter-free and are directly obtained from tokenized BED files. We represent these embeddings as a matrix **B** ∈ ℝ^*V ×N*^, where *N* is the number of tokenized BED files, and *V* is the size of the universe (Figure 1B). If the *l*th region exists in the *n*th tokenized BED file, then the entry at the *l*th row and *n*th column is 1; otherwise the value is 0. The Binary embeddings can be very high-dimensional when the number of tokenized BED files *N* is large. To obtain lower dimensional embeddings, we apply PCA (19) to reduce the dimension of the Binary embeddings from *N* (*N* = 690 in our analysis) to 10 (PCA-10D) or 100 (PCA-100D) (Figure 1B).

#### Region2Vec embeddings

Region2Vec first randomly shuffles regions in each tokenized BED file (Figure 1C), then, similar to training word embeddings in Word2Vec (20), it uses a context window to create prediction tasks and updates Region2Vec embeddings (Figure 1D) (16). Region2Vec embeddings are derived from the first step of region-set2vec (16), which first creates region embeddings, and then averages these to create region set embeddings. Region2Vec maps genomic regions to *D*-dimensional vectors. Region2Vec treats each region as a word, and treats regions in the same tokenized BED file as a sentence. In natural language, words in a sentence often serve to convey a single idea. Similarly, regions in the same BED file share a similar biological context and can be considered as words in a sentence. Then, Region2Vec leverages techniques in natural language processing (NLP) (21) to produce region embeddings. We use the gensim (22) implementation of Word2Vec to implement Region2Vec. There are two steps in the Region2Vec training pipeline: region shuffling and training.

#### Region shuffling

Treating regions in a BED file as a sentence allows Region2Vec to exploit techniques (21) from NLP to produce region embeddings. However, these models are built for natural language, which imposes a specific order between regions. Regions in a BED file do not have any order and can be considered as neighbors of each other since they share the same biological context. To break such order, we randomly shuffle the regions in each region sentence obtained by reading regions in order from a tokenized BED file.

#### Region2Vec training

Region2Vec training uses the region co-occurrence information contained in region sentences to generate region embeddings. Specifically, for each region sentence, Region2Vec moves a context window with length *W* from the beginning to the end of the sentence. At each position of the window, the *W* regions covered by the window are used to update region embeddings via a prediction task. The task asks a Region2Vec model to predict the remaining region given *W* − 1 regions in the context window. We design the Region2Vec model as a three-layer neural network with weights **W**_*I*_ ∈ ℝ^*V ×D*^ and **W**_*O*_ ∈ ℝ^*V ×D*^ for the first and second half of the network, respectively, where *V* is the size of the universe used in tokenization, and *D* is the embedding dimension. The input for the model is a binary vector **a** = [0, …, 1, 0, 1, …, 0] of length *V*. The ones in **a** correspond to *W* − 1 regions in a context window. Then, the model shows how likely each region will be in the context window with a probability distribution **p** ∈ ℝ^*V ×*1^ over all the universe regions. Region2Vec aims to maximize the probability on the remaining region by optimizing **W**_*I*_ and **W**_*O*_. We move the context window and continue the above prediction and optimization procedures until all region sentences have been consumed. The obtained weight matrix **W**_*I*_ are the embeddings for all the universe regions.

#### Region2Vec embeddings training

We consider three important hyperparameters in Region2Vec: context window size *W*, embedding dimension *D*, and initial learning rate *r*. The context window size determines how many region embeddings are jointly updated each time. The embedding dimension specifies the number of vector dimensions used to represent a region. The initial learning rate controls the update magnitude for region embeddings. To explore how these hyperparameters affect the quality of the embeddings, we did a grid search, taking different values of each of these parameters in combination. We take *W* from {5, 50}, *D* from {10, 100}, and *r* from {0.025, 0.1, 0.5} to generate 12 sets of region embeddings for each universe. During Region2Vec training, we shuffle the whole set of tokenized BED files 100 times. After each shuffling, the model is trained on the shuffled files, and after that, the learning rate is linearly decayed until reaching the minimum 10^*−*4^.

For each of these 15 sets of embeddings (Table S1), we compared their performance by calculating two statistical scores: the cluster tendency score (CTS), reconstruction score (RCS), and two biological scores, the genome distance scaling score (GDSS) and neighborhood preserving score (NPS).

### Cluster tendency score (CTS)

The Cluster Tendency Score (CTS) quantifies the degree to which region embeddings can be clustered. Our rationale is that embeddings that form clusters are more likely to be useful than embeddings that are diffuse. We use the idea from the Hopkins test (23) to design our score. Given a set of *N* region embeddings, we first sample *N*_*S*_ embeddings 𝒬 = {**q**_*i*_|*i* = 1, …, *N*_*S*_} from the given set to reduce the computational complexity, where **q**_*i*_ is a region embedding. This step is optional if the time and computational resources can support the evaluation with all *N* region embeddings. We found that using *NS* = 10^4^ is enough to produce stable results.

Then, from the *N*_*S*_ embeddings, we further subsample *N*_*T*_ embed-Dings 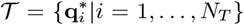 as test points, where 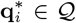. For each test point 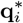, we find its nearest neighbor 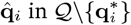, i.e., 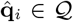 and 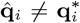. We calculate *D*_*T*_, the summation of all the distances between 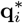 and 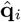, as 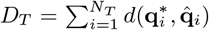. We define the distance function *d*(*·, ·*) as the squared Euclidean distance.

Next, we generate *N*_*T*_ random points 𝒰= {**u**_*i*_|*i* = 1, …, *N*_*T*_}, that are uniformly distributed in the same area where the embeddings in 𝒰 reside. To do this, we uniformly sample **u**_*i*_ between **q**_min_ and **q**_max_, where **q**_min_ and **q**_max_ are vectors. Each dimension of **q**_min_ (**q**_max_) has the minimum (maximum) value of the corresponding dimension of the embeddings in 𝒰. Then, we calculate *D*_*R*_, the summation of all the distances between **u**_*i*_ and its nearest neighbor 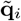 in 𝒬, as 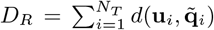. If there are clusters in 𝒬 then the random points should have large distances to their nearest neighbors in 𝒬.

We define the CTS as 2 *·* max(*D*_*R*_*/*(*D*_*R*_ + *D*_*T*_) *−* 0.5, 0) (Figure 2A), a normalized ratio between *D*_*R*_ and *D*_*T*_, ranging from 0 to 1. A larger CTS indicates a large *D*_*R*_ and a greater tendency for the embeddings being evaluated to have clusters. When the embeddings are uniformly distributed, we have *D*_*R*_ *≈ D*_*T*_, *D*_*R*_*/*(*D*_*R*_ + *D*_*T*_) *≈* 0.5, and CTS *≈* 0. Note that points in 𝒰 lie in a hypercube and that the embeddings in 𝒬 may not distribute in that shape in practice. As a result, *D*_*R*_ may be overestimated, inflating the CTS. For example, embeddings in 𝒬 can be uniformly distributed in a sphere within the hypercube, and sampling points in 𝒰 at the corners of the hypercube will increase *D*_*R*_, leading to a large CTS even when there are no clusters in 𝒬. We address this by shrinking the hyper-cube a little bit, using 95% percentile and 5% percentile of each dimension data in 𝒬 to define **q**_max_ and **q**_min_, respectively. If the embeddings being evaluated deviate from randomly distributed ones, then *D*_*R*_ will be large, and *D*_*T*_ will be relatively small; therefore, their CTS will be large.

**Figure 2.**
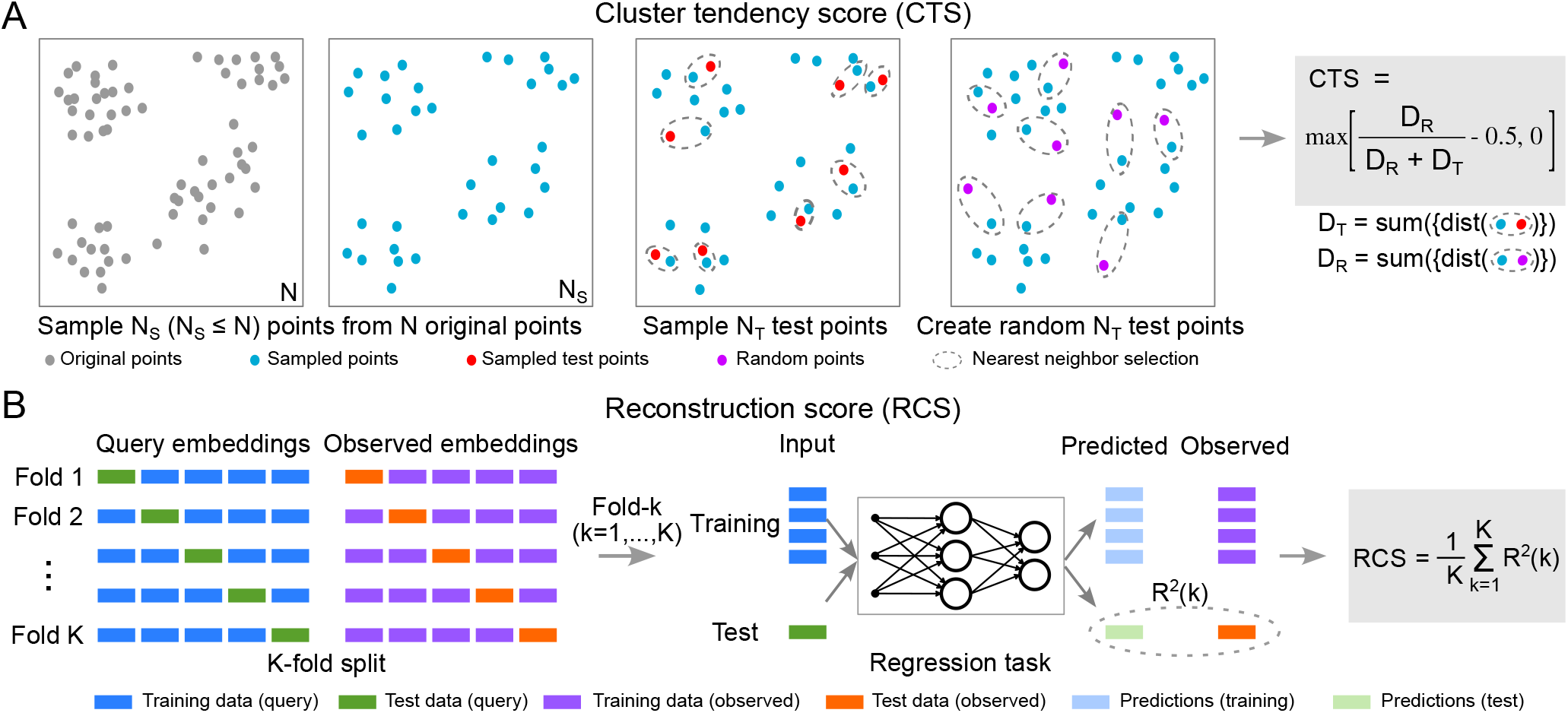
Overview of the two statistical evaluation scores. **A**. Illustration of calculating CTS. **B**. Illustration of calculating RCS.

### Reconstruction score (RCS)

One limitation of CTS is that it evaluates only the spatial distribution of region embeddings, and does not assess the information contained in the embeddings. To address this, we sought a score that could evaluate embedding content more directly. To this end, we developed the Reconstruction Score (RCS), a statistically motivated metric that measures how much embeddings preserve the information about regions’ occurrence among the training BED files. To measure how much such information is preserved in a region embedding, we create a regression task to predict the original input vectors, and evaluate the performance of region embeddings on this task. Intuitively, RCS measures how well the embeddings can be used to reconstruct the full-dimensional input data.

To compute RCS, we first define the Binary embeddings as “observed” embeddings, since they represent all the regions’ occurrences in the training data by definition. These Binary embeddings **B** (row vectors) are the labels in the regression task. We define the trained embeddings as “query” embeddings **Q** (row vectors), which are the inputs in this task. Each region thus has a “query” and an “observed” representation. We preprocessed both the query and observed embeddings so that they have zero mean and unit variance. We split **B** and **Q** into *K* folds (Figure 2B, K-fold split) **B** = [**B**_1_, …, **B**_*K*_ ] and **Q** = [**Q**_1_, …, **Q**_*K*_ ]. Then, we use each fold to train a regression model *f*_*θ*_ with parameters *θ* that predicts the observed embeddings **B**_*k*_ using the corresponding query embeddings **Q**_*k*_ (Figure 2B, Regression task). We instantiate *f*_*θ*_ as a three-layer feed-forward neural network with 200 hidden units and ReLU activations. The input dimension of the model depends on the dimension of given region embeddings *D*, and the output dimension of the model is the number of BED files in the collection (*N* = 690). The performance of *f*_*θ*_ reflects quality of **Q** in this evaluation. We use mean-squared error as the training loss. During evaluation, we use *R*^2^ score because it is bounded between 0 and 1 and leads to an interpretable score with 1 indicating perfect reconstruction capability. We calculate *R*^2^(*k*) score for the *k*th fold defined as follows

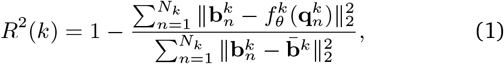

where *N*_*K*_ denotes the number of rows in 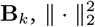 denotes the squared 2-norm of a vector, 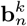 denotes the *n*th row of **B**_*k*_, 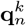 denotes the *n*th row of **Q**_*k*_, 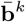 denotes the row average of **B**_*k*_, and *θ*^*k*^ denotes the parameters learned at the *k*th step. We repeat the above process process for *K* steps. Finally, the RCS for **Q** is the average *R*^2^ score over the *K*-fold cross-validation, i.e., 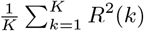. To make the RCS for different sets of embeddings comparable, we keep the architecture of the neural network *f*_*θ*_ fixed.

In theory, Binary embeddings should have the highest RCS of 1; however, the model cannot perfectly be trained to be an identity function, so we consider the score of the Binary embeddings to be an upper bound. We used the implementation from scikit-learn (24) and calculated the RCS of the region embeddings that we generated for the seven universes. To reduce the time and space complexity induced from a large *N*, we randomly sample a small portion of the training BED files, e.g., 10 files, for evaluation. We demonstrated in the Results section that this technique does not significantly affect the evaluation between different sets of region embeddings.

### Genome distance scaling score (GDSS)

CTS and RCS provide useful insight into the clusterability and reconstruction ability of embeddings; however, neither of them considers biological knowledge. Thus, we next sought to devise evaluations that take into account the biology. To this end, we developed the Genome Distance Scaling Score (GDSS), which quantifies how much the embedding distances vary with the corresponding genome distances for a set of region embeddings. The GDSS analyzes how the embedding distance (ED) between two regions changes with their genomic distance (GD). The ED and GD reflect two different relationships between two regions; the ED reflects their region co-occurrence in BED files, whereas the GD reflects their distance in genome space. Since regions that are close on the genome tend to have more similar biological functions than regions that are distant on the genome, we reasoned that, on average, the GD could be used as a noisy proxy for *some* shared biological function. Of course, regions that are distant from one another on the genome may be functionally similar, and may also be near one another in embedding space because of their functional similarity in biology. However, on average, we expect GD to be smaller between two regions that are similar than between two random regions. In training the embeddings, the location on the genome is not provided to the model. Therefore, any correlation between genomic distance and embedding distance reflects biological learning accomplished by the training process. We exploit this expectation to design the GDSS, reasoning that embeddings that capture this biological signal will have GD positively correlated with ED.

Therefore, to calculate GDSS, we first randomly sample *N*_*S*_ region pairs from the universe and calculate their ED and GD values (Figure 3A). The GD between two regions 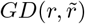 is calculated as follows,

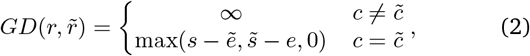

where *r* = (*c, s, e*) 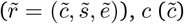 represents the chromosome index of 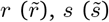 and 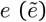 denote the start and end positions of 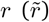, respectively. Therefore, if two regions overlap, the GD is zero; if they are on different chromosomes, the GD is infinity; otherwise, the GD is the smallest number of base pairs connecting the two regions. The ED between the two regions 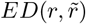 is the cosine distance between their region embeddings because the distance is immune to the artifacts caused by the magnitudes of region embeddings or the embedding dimensions. For example, the Euclidean distance between two region embeddings obtained from Region2Vec can be simply enlarged using a large learning rate or a large embedding dimension, leading to a large GDSS. However, using cosine distance does not have this issue since the distance is bounded from -1 to 1.

**Figure 3.**
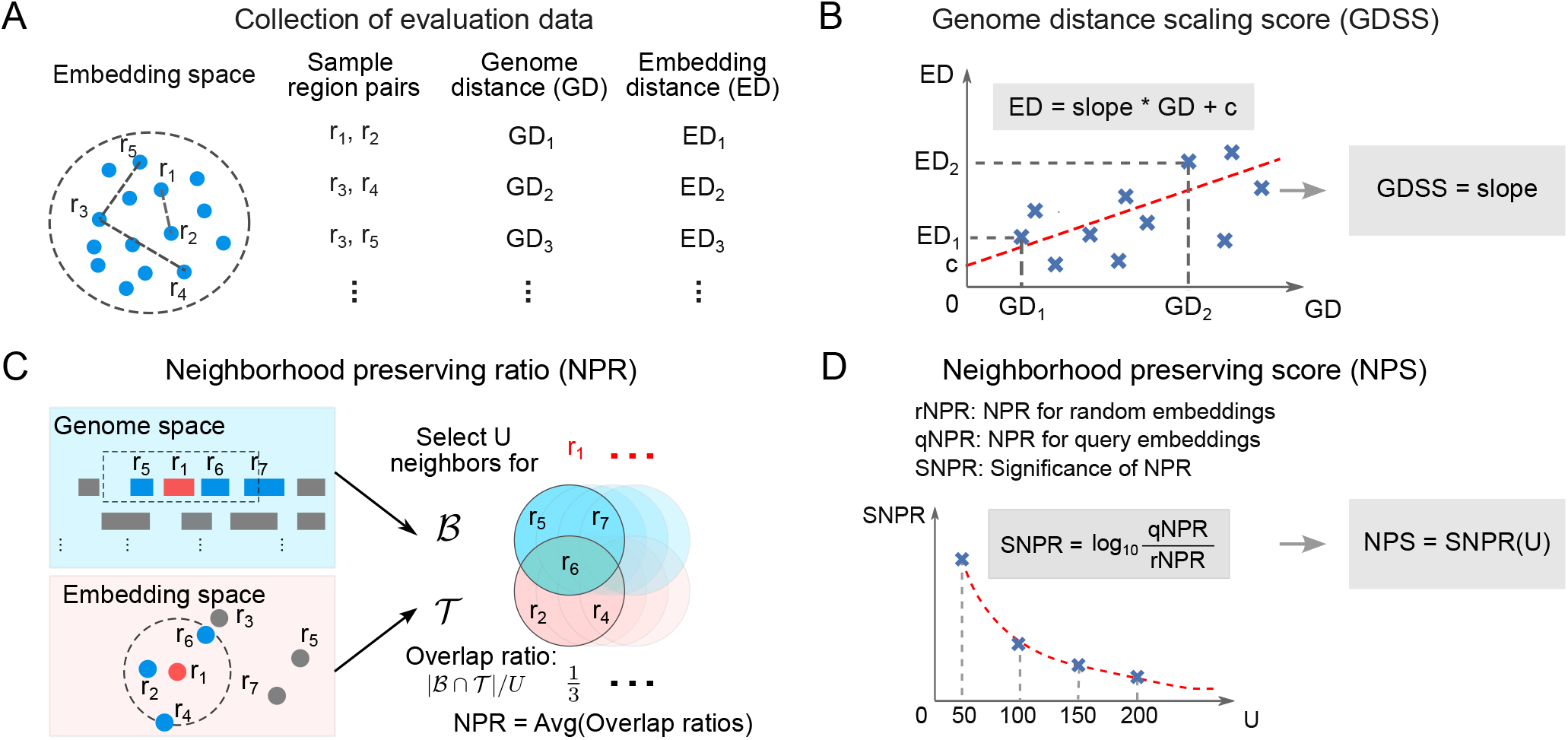
Overview of the two biological scores. **A**. GDSS calculation Step 1: Sample region pairs from a universe and calculate ED and GD for each pair. **B**. GDSS calculation Step 2: Calculate the slope of the linear curve fitted on (ED, GD) pairs as the GDSS. **C**. NPS calculation Step 1: Calculate NPR with a neighborhood size U as the average of the overlap ratios for all sampled regions. **D**. NPS calculation Step 2: Calculate how significant the NPR of a set of query embeddings given U is over a set of random embeddings as the NPS.

Finally, we conduct a linear curve fitting for the *N*_*S*_ ED and GD pairs and obtain the slope of the fitted curve as the GDSS for the set of region embeddings being evaluated (Figure 3B). A positive score indicates the set of region embeddings captures the biological information that the closeness of regions in the genome reflects their functional similarity. The magnitude of the score indicates how strong such information is captured by the region embeddings. A large GDSS is often associated with embeddings with large and low-density clusters, where the embeddings tend to repel each other to have a large average ED.

GDSS is not a perfect score of biological information capture. In fact, a model that is provided with genomic locations in the training process could exploit the locations to place region embeddings near one another when the regions are near in genome space; however, such a model will not have learned anything useful. Nevertheless, for a model that is *not* provided with genomic locations in training, the ability to recapitulate genome distance to some degree provides evidence that the model has learned underlying biology.

### Neighborhood preserving score (NPS)

GDSS focuses on individual region pair distances and does not consider local neighborhood structures. We thus developed the Neighborhood Preserving Score (NPS), which finds whether the regions neighboring on the genome are also close in the embedding space. Like GDSS, NPS relies on the general tendency of biological regions to cluster by function. To achieve this, NPS considers the overlap between a region’s neighboring set in the genome space and the corresponding set in the embedding space.

NPS quantifies the degree to which linear genomic neighborhood information of genomic regions is preserved in the embedding space. Compared with GDSS, NPS does not evaluate individual region embeddings; instead, it focuses on local sets of region embeddings. It is based on the same assumption that a region’s neighboring regions in the genome space tend to have similar biological functions, and whether the embeddings capture this relationships can be used as a measure of their quality. Hence, for good embeddings, we expect to observe a significant overlap between the neighboring region sets from the genome and the embedding space.

NPS uses SNPR to measure the significance of the overlap in average. Specifically, we first calculate the overlap between the *U* - nearest neighbors of a region in genome space, ℬ^*U*^ (*r*) and its *U* - nearest neighbors in embeddings space, 𝒯^*U*^ (*r*) of a region *r*. We use the GD defined in Eq. (2) and cosine distance as the ED. Then, we calculate the overlap ratio *ρ*(*r*) for the region *r* as follows:

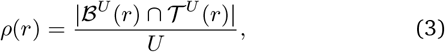

where | *·* | denotes the size of a set. Averaging over all *ρ*(*r*) gives us the NPR (Figure 3C) for the set of region embeddings being evaluated. We denote such an NPR as qNPR. To measure how significant the NPR is, we calculate the NPR for a set of random region embeddings, denoted as rNPR. A random region embedding is created by sampling a value from -0.5 to 0.5 for each embedding dimension. Finally, we calculate SNPR as

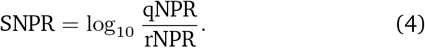

We calculate SNPR (Figure 3D) as the NPS for a set of region embeddings. A large NPS indicates the strong capability of the region embeddings to preserve the neighborhood information. When the value of NPS is close to 0, the *U* -neighborhood preserving capability of the region embeddings is only as good as a set of random embeddings. Increasing *U* increases the distance of the regions on the genome and decreases the value of qNPR. Hence, NPS will also decrease. In contrast, a very small *U* may lead to zero qNPR, making NPS an ineffective measure of evaluating region embeddings. In our evaluation, we set *U* as a multiple of 50 so that we can effectively compare different sets of region embeddings using NPS.

### *F*_10_ score

Finally, in the process of computing and evaluating these scores, we realized that the quality of embeddings may be affected by the goodness of fit of the universe. We therefore needed a way to assess the universe fit to provide more insight into how the scores perform. In parallel work, we recently developed the *F*_10_ score as a way to assess fit of a universe to a collection (25). Briefly, the *F*_10_ score is computed as a weighted harmonic mean of the precision and recall when considering a query region set as a positive file with a universe as a prediction of that file. Specifically, we first consider a genomic position covered by a raw region set in the collection and ask whether this position covered by the universe. We count the number of genomic positions that are covered in both as true positive (TP), the number covered only in the universe as false positive (FP), and the number covered only in the region set as false negative (FN). With TP, FP, and FN, we calculate precision and recall. Precision tells us the ratio of the universe covering the region set, and recall gives the ratio of the region set covering the universe. We then compute the *F*_10_ score, which is similar to the *F* score, and combines the precision and the recall with the harmonic mean, except we weigh the recall 10 times more weight than precision, due to the asymmetry of the relationship between universe and region set collection member. The asymmetry of the weights favors a universe that broader than the region set, since the universe needs to be more general to accommodate many different region sets. The *F*_10_ score ranges between 0 and 1, with 1 representing the perfect fit between the universe and the region set. We applied the this score to each embedding set to improve interpretability of the evaluation metrics.

## Results

### Cluster tendency score (CTS)

To calculate CTS, we first sample *N*_*S*_ = min(10^4^, *N*) region embeddings from the original *N* embeddings to reduce the computational complexity. Then, from the *N*_*S*_ embeddings, we sample *N*_*T*_ = 10% *· N*_*S*_ test points. For each of the *N*_*T*_ test points, we compute the distance to its nearest neighbor among the *N*_*S*_ sampled points excluding itself. We get the distance summation over the *N*_*T*_ test points as *D*_*T*_. We do the same for *N*_*T*_ random points and get the distance summation as *D*_*R*_. Next, we calculate the CTS for each set of region embeddings. To combat the randomness in the score, we computed CTS with 20 different random seeds and report the average.

We observed the embeddings being evaluated tend to cluster with a large CTS (e.g., Figure 4A). In contrast, a lower CTS indicates the embeddings are more dispersed throughout the embedding space (e.g., Figure 4B), with a score near 0 indicating no clusters (e.g., Figure 4C). We also observed that CTS varied across embedding approaches, and that CTS is robust to the choice of universes (Figure S1). Scores for the 5k tiling universe varied from 0.84 (5W100D-0.0250r) to 0.00 (5W10D-0.5000r) (Figure 4D). Interestingly, the PCA-10D embeddings have a higher score than the PCA-100D and Binary (Figure 4D), even though the PCA-10D retains less information from the training data than the PCA-100D and Binary. In other words, a high CTS does not necessarily indicate better embeddings, since we can always generate a set of region embeddings with a very high score while being irrelevant to the training data. Instead, a CTS is only relevant to the spatial distribution of region embeddings and quantifies how likely they can be clustered. For Region2Vec embeddings, we observed that small context windows (e.g., *W* = 5) achieved higher CTSs than large context windows (e.g., *W* = 50). Moreover, using a large initial learning rate (e.g., *r* = 0.5) leads to training failures since the corresponding CTSs are around 0, indicating that the embeddings are just as good as random embeddings in terms of clusterability. Therefore, for training Region2Vec embeddings with good clusterability, avoid using a large *r*, and use a small *W*. We conclude that the CTS is useful to identify low quality region embeddings whose CTS is near 0.

**Figure 4.**
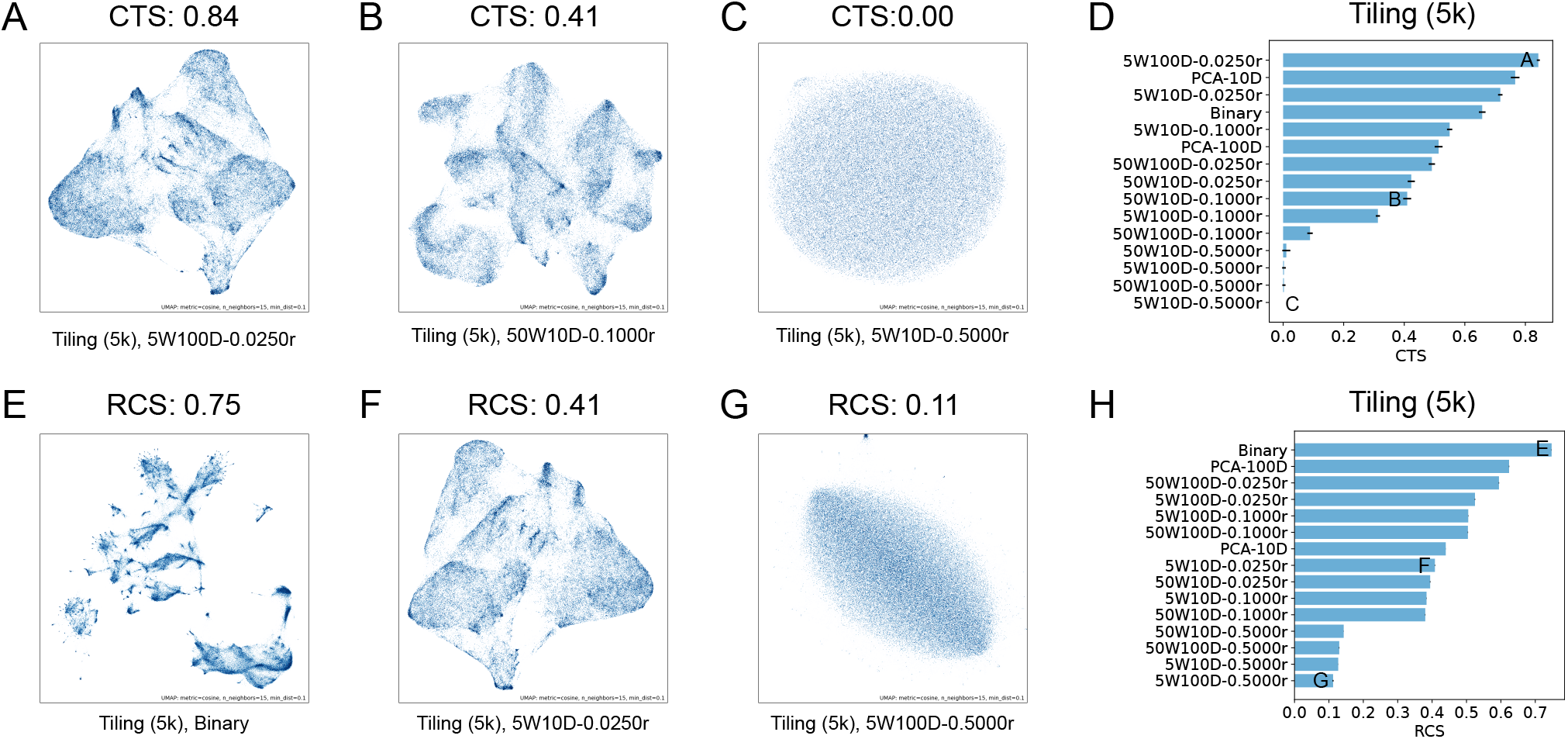
CTS and RCS results for the Tiling (5k) universe. **A**-**C**. UMAP visualizations of the 5W100D-0.025r, 50W100D-0.1000r, and 5W10D-0.5000r embeddings. **D**. CTSs of different sets of region embeddings. **E**-**G**. UMAP visualizations of the Binary, 5W10D-0.0250r, and 5W100D-0.5000r embeddings. **H**. RCSs of different sets of region embeddings. Each blue horizontal bar and the associated black bar in **D** (**H**) indicate the average and standard deviation of the CTSs (RCSs) from 20 (5) runs of the CTS (RCS) calculation for each set of region embeddings. W: context window size, D: embedding dimension, r: initial learning rate.

### Reconstruction score (RCS)

We used 5-fold cross validation to calculate RCSs. Binary embeddings faithfully represent the original similarity information between regions in the training data and demonstrate that the data has rich structures (Figure 4E). For Region2Vec embeddings learned with different context window sizes, a high RCS implies a large context window size and rich structures in the corresponding embeddings (e.g., Figure 4F). We reason that a large context window can well preserve the region occurrence information contained in the training data, since in theory, all regions in a BED file should be in the same context. Thus, Region2Vec embeddings learned in this way capture various degrees of similarity between genomic regions, exhibiting rich structures in the embedding space. In contrast, using a small context window tends to move regions in the same BED file but not in the same context window away from each other, perturbing the original similarity information between regions. Region2Vec embeddings learned with a smaller context window tend to have a lower RCS and exhibit less distinctive structures (e.g., Figure 4G).

Computing RCSs across our embedding sets, we observed that RCSs are generally robust to the choice of universe (Figure S2). Specifically, on the 5k tiling universe, as expected, Binary embeddings achieve the highest RCS (Figure 4H). The PCA embeddings have decreasing RCSs as dimensionality is decreased, consistent with fewer dimensions retaining less information about the original data (Figure 4H). Among the Region2Vec embeddings, across universes, 50W100D-0.0250r achieves the highest RCS and 5W100D-0.5000r achieves the lowest; in general, using a large context window (e.g., *W* = 50), a large embedding dimension (e.g., *D* = 100), and a small initial learning rate (e.g., *r* = 0.025) scores higher, indicating that these training settings best preserve the region occurrence information contained in the training data (Figure 4H).

Compared with CTS, which favors a small context window (*W* = 5), RCS favors a large context window (*W* = 50), which demonstrates a tradeoff between the clusterability of Region2Vec embeddings and their capability in preserving the region occurrence information contained in the training data. If we prioritize clusterability, we should choose a small context window, whereas if we prioritize preserving information for downstream data processing, we should choose a larger context window.

The space and time complexities of calculating RCS increase when the number of output dimensions or the number of training BED files *N* increases. To address, we randomly sample a small portion of the given BED files for the RCS calculation. We experimented with 690 (all) BED files and 10 BED files and calculated the Pearson correlation coefficient between the RCSs for the 15 sets of region embeddings under each universe. We observe that reducing the number of output dimensions does not significantly affect the relative measurements between different sets of region embeddings under each universe (Table 1).

**Table 1.**
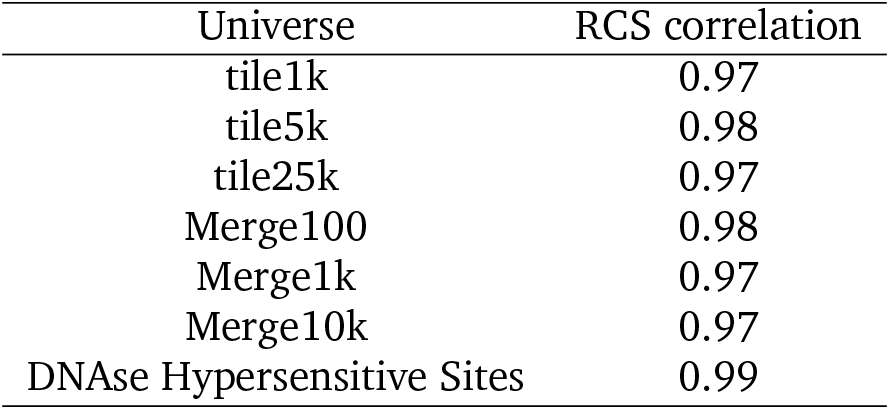
Pearson correlation coefficients between RCSs calculated with 690 and 10 output dimensions for region embeddings under each universe.

### Genome distance scaling score (GDSS)

To calculate GDSS, we first randomly sample 100k pairs of regions and calculate the ED and the GD for each region pair. Then, we compute the line of best fit over the 100k (GD, ED) points. The GDSS is the slope of the fitted line showing the global trend of ED as GD increases.

We first show how the GDSS varies visually. We observed that GDSS is higher for embeddings with low-density clusters. For example, we can observe large and sparse clusters from the high-scoring embeddings 50W10D-0.0250r for Merge (100) (Figure 5A) and PCA-10D for Merge (10k) (Figure 5E). In contrast, we can observe dense clusters from the low-scoring embeddings Binary for Merge (100) (Figure 5B), PCA-10D for Merge (100) (Figure 5C), and Binary for Merge (10k) (Figure 5F). For almost random embeddings, there is no clear cluster (e.g., Figure 5G).

**Figure 5.**
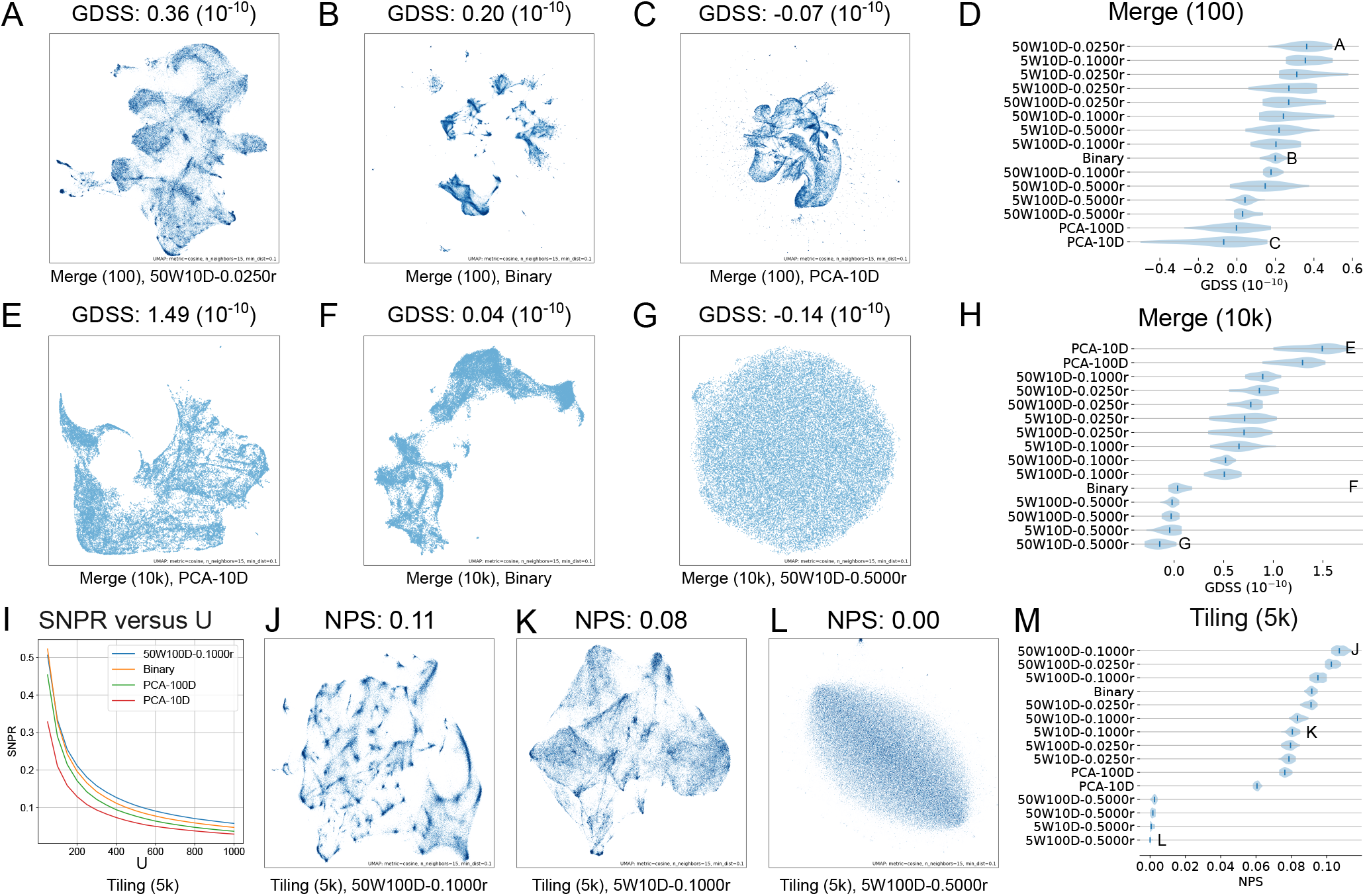
GDSS and NPS results. **A**-**C**. UMAP visualizations of the 50W10D-0.0250r, Binary, and PCA-10D embeddings for the Merge (100) universe. **D**. GDSSs for different sets of region embeddings for the Merge (100) universe. **E**-**G**. UMAP visualizations of the PCA-10D, Binary, and 50W10D-0.5000r embeddings for the Merge (10k) universe. **H**. GDSSs for different sets of region embeddings for the Merge (10k) universe. **I**. SNPR versus U curves for the selected embeddings for the Tiling (5k) universe. **J**-**L**. UMAP visualizations of the 50W100D-0.1000r, 5W10D-0.1000r, and 5W100D-0.5000r embeddings for regions in the Tiling (5k) universe. **M**. Calculate NPS with U = 500 for different sets of region embeddings of the seven universes, respectively. Each shaded area and blue vertical bar in **D, H**, and **M** indicate the distribution and median of GDSSs/NPSs over 20 runs of the calculation for each set of embeddings. W: context window size, D: embedding dimension, r: initial learning rate.

Computing the GDSS for each set of embeddings in Table S1 across universes, we observed that unlike the statistical evaluations, the GDSS rankings vary substantially by universe. For example, in the Merge (100) universe, the PCA embeddings have the lowest GDSSs (Figure 5D), whereas, for the Merge (10k) universe, the PCA embeddings have the highest GDSSs (Figure 5H).

We next sought to investigate why the universe selection has such a drastic effect on GDSS. We reasoned that the tokenization under a universe could mask the shared function of nearby regulatory elements. For example, after tokenization, nearby regions are merged into large consensus regions, or they are completely missed due to the lack of coverage of the universe (25). Then, the shared function signature could disappear, leading to lower GDSSs. To test this, we sought to determine the “fit” of each universe to the input data. We computed the *F*_10_ scores (see Methods), a score we developed recently (25) to measure how well a chosen universe covers the regions in the provided BED files (before tokenization). Larger *F*_10_ scores indicate the universe is a better fit for the regions in the collection. Computing *F*_10_ on the seven universes, we found that the Merge (100) universe has a higher *F*_10_ score (*F*_10_ = 0.6281) than the Merge (10k) universe (*F*_10_ = 0.1990), indicating it is a better fit to the input collection. This may explain why the embeddings tokenized with different universes have such different rankings; using the Merge (100) universe for tokenization can preserve more information contained in raw BED files and is less noisy than tokenizing with Merge (10k), for which simply using PCA to reduce noise works better than using Region2Vec to learn from very noisy data. Similarly, Tilling (25k) has a low *F*_10_ score (*F*_10_ = 0.1591), and PCA embeddings perform the best in terms of GDSS (Figure S3F).

Interestingly, Region2Vec embeddings perform best when the universe is a good fit to the BED files (Figure S3A,B,D,E,G). This indicates that Region2Vec is capturing biological knowledge in the embeddings. We conclude that using a universe with a high *F*_10_ score is important for Region2Vec. The context window size *W* depends on the size of the universe. Using *W* = 5 is preferred for a large universe (Figure S3A,D,G), while using *W* = 50 is preferred for a relatively small universe (Figure S3B,C,E,F). Avoid using a very large initial learning rate (e.g., *r* = 0.5) since it leads to very low GDSSs. In most cases, using a small embedding dimension (e.g., *D* = 10) is beneficial for achieving a high GDSS.

### Neighborhood preserving score (NPS)

To extended GDSS beyond individual region pair distances and consider local neighborhood structure, we next computed the NPS. We randomly sampled 10^4^ regions, and for each sampled region, we obtain its sets of *U* -nearest neighbor regions in the genome space (*B*) and its *U* -nearest neighbor regions in the embedding space (*T*). Then, we computed NPSs for each of the 15 sets of generated region embeddings for each universe.

As *U* increases, we observe decrease in NPSs (Figure 5I), which is expected since the sharing in biological function decreases as the neighboring regions get further away in genome space as *U* increases. For comparison using NPSs, we choose a large *U*, i.e., *U* = 500, since we are more interested in testing whether a set of region embeddings captures rich biological information which is often contained in a large neighborhood of regions. Although we choose *U* = 500 in Figure 5J-M, the ranks of different sets of region embeddings are stable under different *U* s (Supplemental Figure S5A-G). An exception happens when *U* is small. For example, when *U* = 50, among different sets of region embeddings for the Merge (100) universe, Binary has the highest NPS (Supplemental Figure S5A).

To achieve a high NPS, the region’s neighboring regions in the embedding space should be as close as possible so that they will not be included in other region’s neighborhood set and decrease the overlap ratio for that region. Therefore, for a set of region embeddings with a high NPS, we will observe many small and dense clusters. For example, we can observe small and dense clusters from the high-scoring embeddings 50W100D-0.1000r for Tiling (5k) (Figure 5J). We observe relatively sparse clusters for embeddings with smaller NPSs (Figure 5K and L). Among different kinds of Region2Vec embeddings, the ones trained with the largest initial learning rate *r* = 0.5 have very small NPSs, and the ones with the large context window size *W* = 50 and the large embedding dimension *D* = 100 have large NPSs (Figure 5M). This observation is consistent across the seven universes (Figure S4). Moreover, for each of the seven universes, Binary, PCA-100D, and PCA-10D have decreasing NPSs. As in other evaluations, using large initial learning rate (*r* = 0.5) leads to ineffective training of region embeddings and small NPSs. High NPSs are best achieved with a large context window (e.g., *W* = 50), a large embedding dimension (e.g., *D* = 100), and a small initial learning rate (e.g., *r* = 0.025).

### Downstream task performance

To demonstrate that our proposed metrics are indicative of the performance of downstream tasks, we used the metadata of the BED files and designed two downstream classification tasks: antibody type classification and cell type classification. We used 60% of the BED files as the training data and the remaining ones as the test data. We first calculated the average embedding of all the tokenized region embeddings in a BED file as the vector representation for the BED file. Then, we trained an SVM classifier with a linear kernel to predict the antibody type or the cell type of the BED file given its vector representation. We used five-fold cross-validation with the F1 score with micro average to measure the classification performance. For the two particular classification tasks, we observe that all the scores have positive correlations with the downstream classification performance, and NPS is the most indicative metric (Table 2). However, there is clear varying in performance;for example, CTS does not predict performance particularly well on these two classification tasks. Nevertheless, as the four proposed scores evaluate different perspectives of region embeddings, choosing the most indicative score depends on the downstream task.

**Table 2.** Pearson correlation coefficients between the classification performance and the four evaluation scores.

| Class type | CTS | RCS | GDSS | NPS |
| --- | --- | --- | --- | --- |
| Antibody | 0.14 | 0.76 | 0.52 | 0.88 |
| Cell | 0.22 | 0.78 | 0.48 | 0.88 |

## Discussion and conclusion

In this paper, we proposed two statistical-based scores, CTS and RCS, and two biological-based scores, GDSS and NPS, to evaluate the quality of genomic region embeddings learned from unsupervised methods. Each score evaluates different perspectives of region embeddings. CTS quantifies how well region embeddings can be clustered, and RCS measures how much original training information is preserved by region embeddings. GDSS calculates the degree to which region embedding distances scale with genomic region distance in the embedding space, while NPS calculates how well region embeddings preserve a region’s neighborhood in genome. These four scores provide complimentary perspectives that, when taken together, can provide a comprehensive evaluation of the statistical and biological value of region embeddings in an unsupervised way. The scores also vary in terms of input requirements: calculating a CTS only needs region embeddings for evaluation, whereas calculating a RCS also needs access to the training data (the tokenized BED files). The biological GDSS and NPS do not need the training data, but they do need the genomic coordinates of regions in order to calculate genome distance.

The four scores do not necessarily match since they evaluate different perspectives of region embeddings. Choosing a good score depends on the task that we focus on. For example, RCS favors Region2Vec embeddings with a large context window, while CTS favors Region2Vec embeddings with a small one. When the task is to identify clusters, it is better to select a set of region embeddings with a high CTS. However, if the the task is to distinguish between different cell/antibody types with high accuracy, it is better to select a set of region embeddings with a high RCS. Moreover, we have found the four scores useful in tandem to ensure region embeddings meets our needs. For example, we can first calculate RCSs of several candidate sets of region embeddings to filter out low-scoring sets that lose too much training information. Then, to ensure biological signatures are preserved, we can calculate GDSS or NPS to further select region embedding sets. Finally, we can calculate CTS to find an easily-clustered region embedding set to facilitate downstream analyses.

Scores of the same type are not directly comparable across universes, since different universes have different sets of regions, resulting in different training data across universes. However, all the scores, except the GDSS, yielded similar rankings regardless of universe. In fact, GDSS is the only one that was sensitive to the choice of universe. Although evaluating the quality of a universe is not our goal in this paper, we found a correlation between GDSS and *F*_10_ score, a measure of universe quality. Therefore, we propose that GDSS may be useful as a way to determine whether a universe is a poor fit, which coincides to the case when PCA embeddings outperform Binary embeddings, providing a way to assess a universe without the need to access the original BED files (before tokenization).

Overall, based on the results from the four scores, we found that with proper settings, embeddings learned from Region2Vec, are top performers in general. More specifically, the Region2Vec embeddings outperformed the binary and PCA embeddings in terms of CTS, GDSS and NPS. We notice that Region2Vec embeddings had lower RCSs than PCA embeddings. We reason that Region2Vec exploits regions’ co-occurrence information in the training data for dimensionality reduction with a finite-sized context window. It may lose certain region co-occurrence information contained in a long region sequence. Compared to Region2Vec, PCA performs relatively simple linear dimensionality reduction on the original data.

Therefore, it is easier for a neural network to learn an inverse function for PCA embeddings to reconstruct the original data (hence, higher RCSs) than for Region2Vec embeddings. Nevertheless, Region2Vec embeddings have demonstrated the ability to capture the simple biology that close regions on the genome have similar biological functions. Our scores also provide parameter selection guidelines for learning good Reigon2Vec embeddings. We expect that these scores will provide a useful basis for evaluation of unsupervised region embeddings, and also anticipate that they can form the basis of evaluation approaches for upcoming supervised region embedding methods as well (26).

## Supplementary information

**Table S1.**
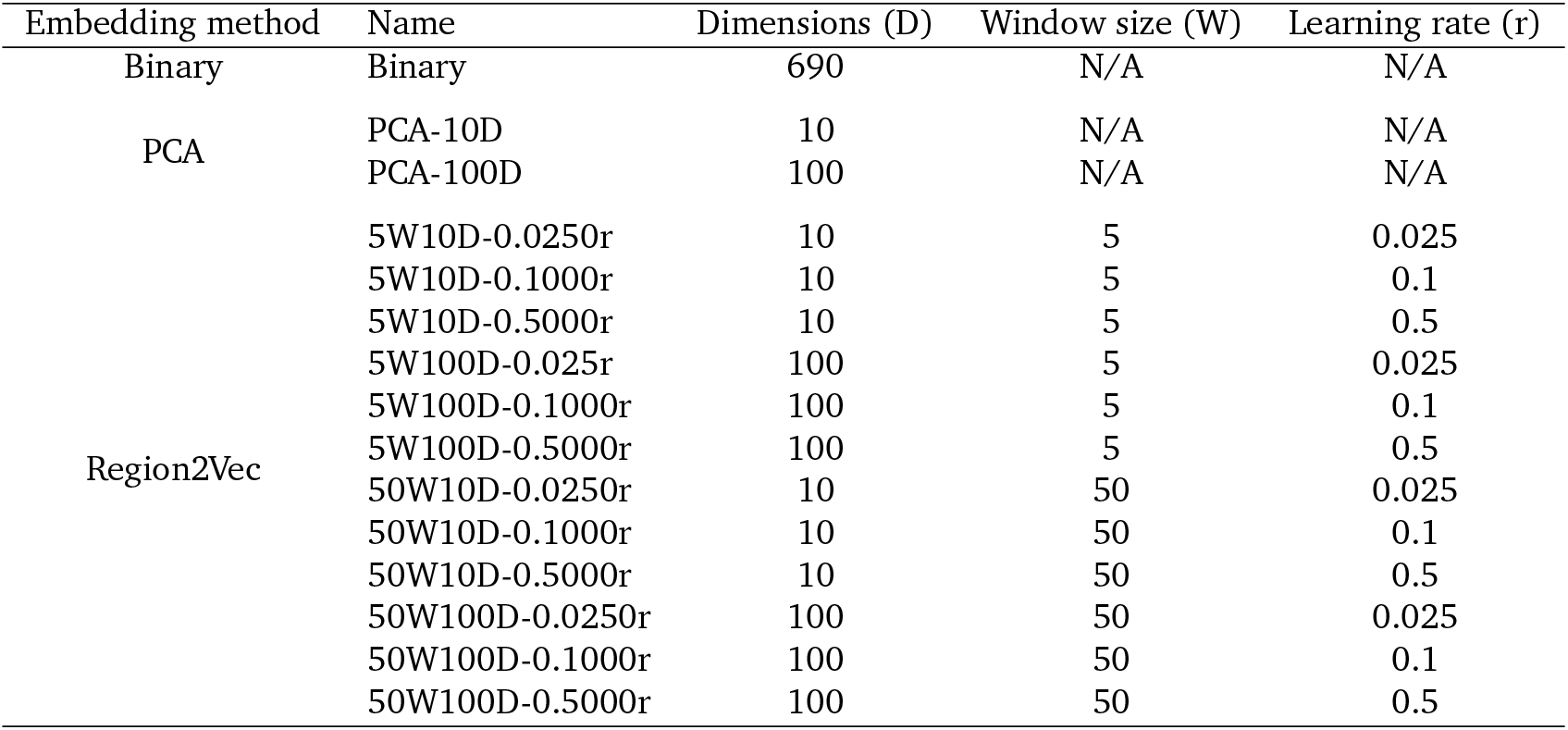
Configurations of the 15 types of embeddings.

**Table S2.**
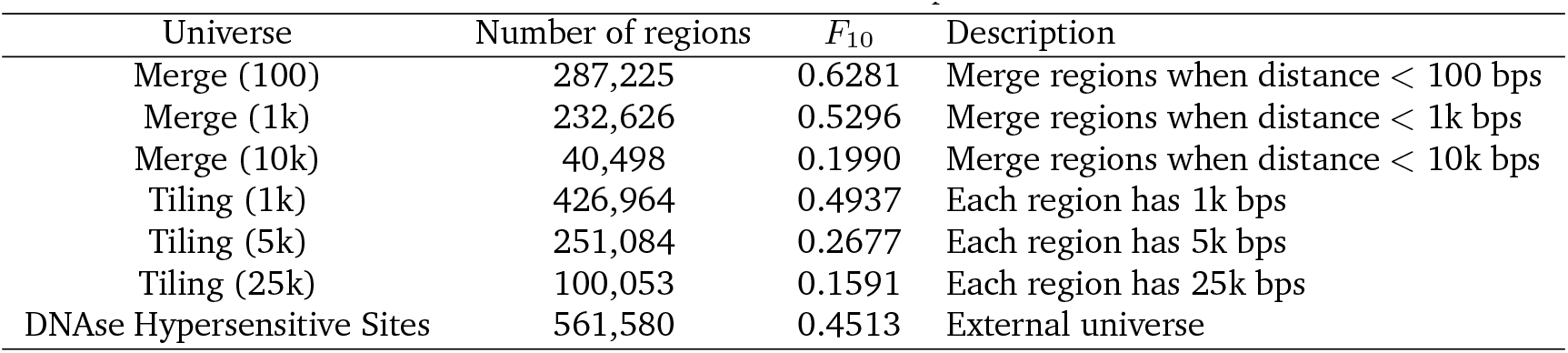
Universes used in the experiments.

**Figure S1.**
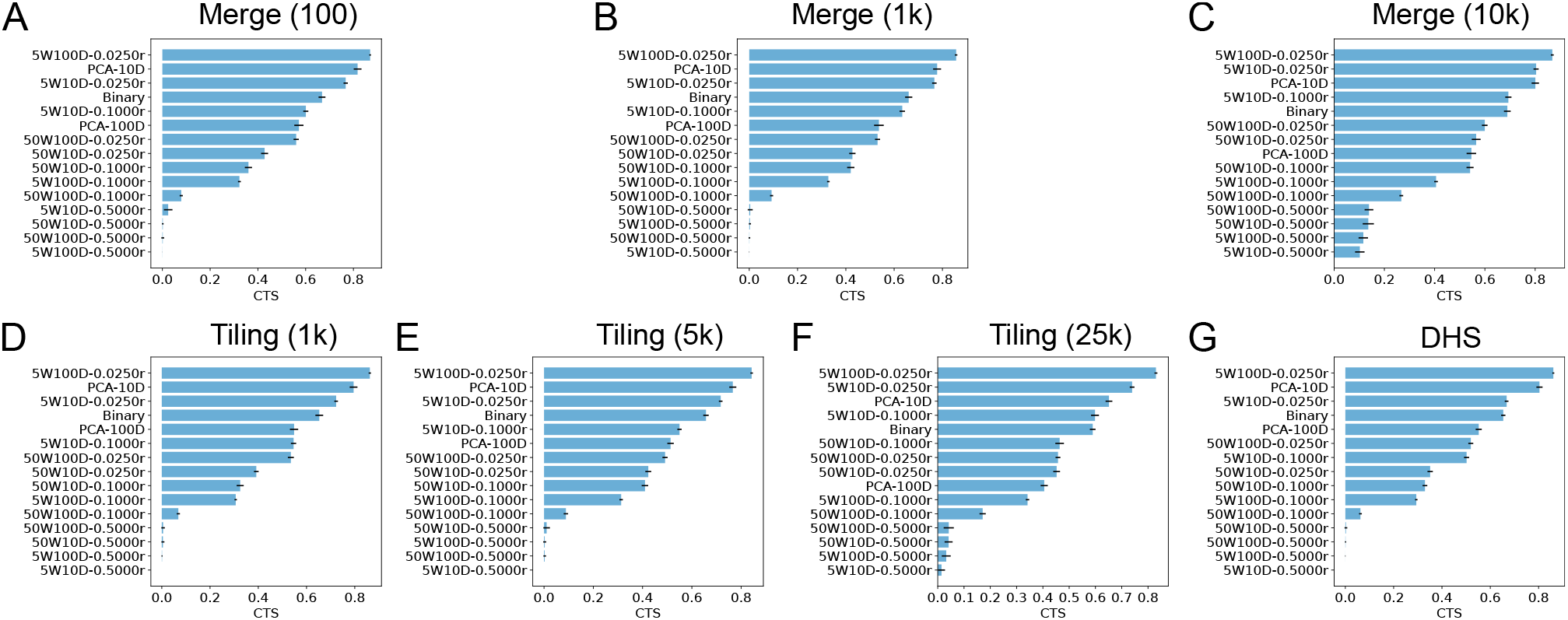
CTS scores of different sets of embeddings for regions in the seven universes. Each blue horizontal bar and the associated black bar indicate the average and standard deviation of the CTSs from 20 runs of the CTS calculation for each set of region embeddings. W: context window size, D: embedding dimension, r: initial learning rate.

**Figure S2.**
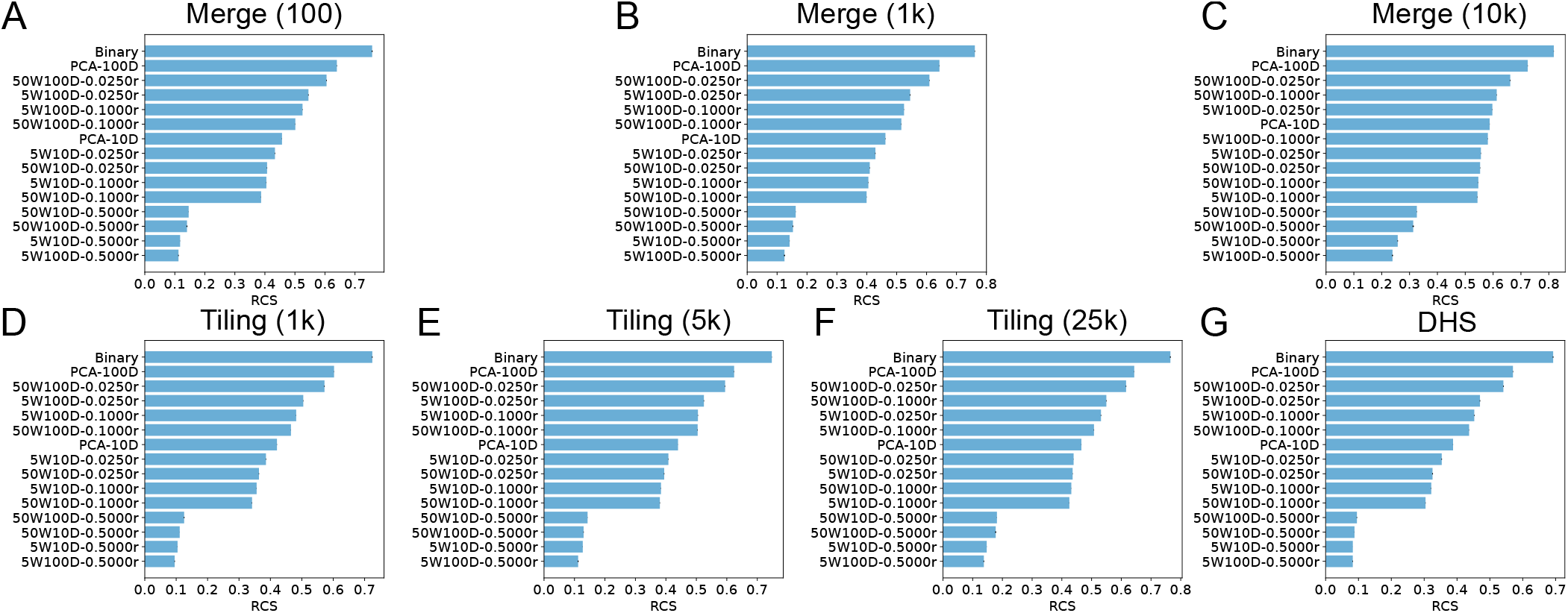
RCSs of different sets of embeddings for regions in the seven universes. Each blue horizontal bar and the associated black bar indicate the average and standard deviation of the RCSs calculated with 5 different random seeds for each set of region embeddings. W: context window size, D: embedding dimension, r: initial learning rate.

**Figure S3.**
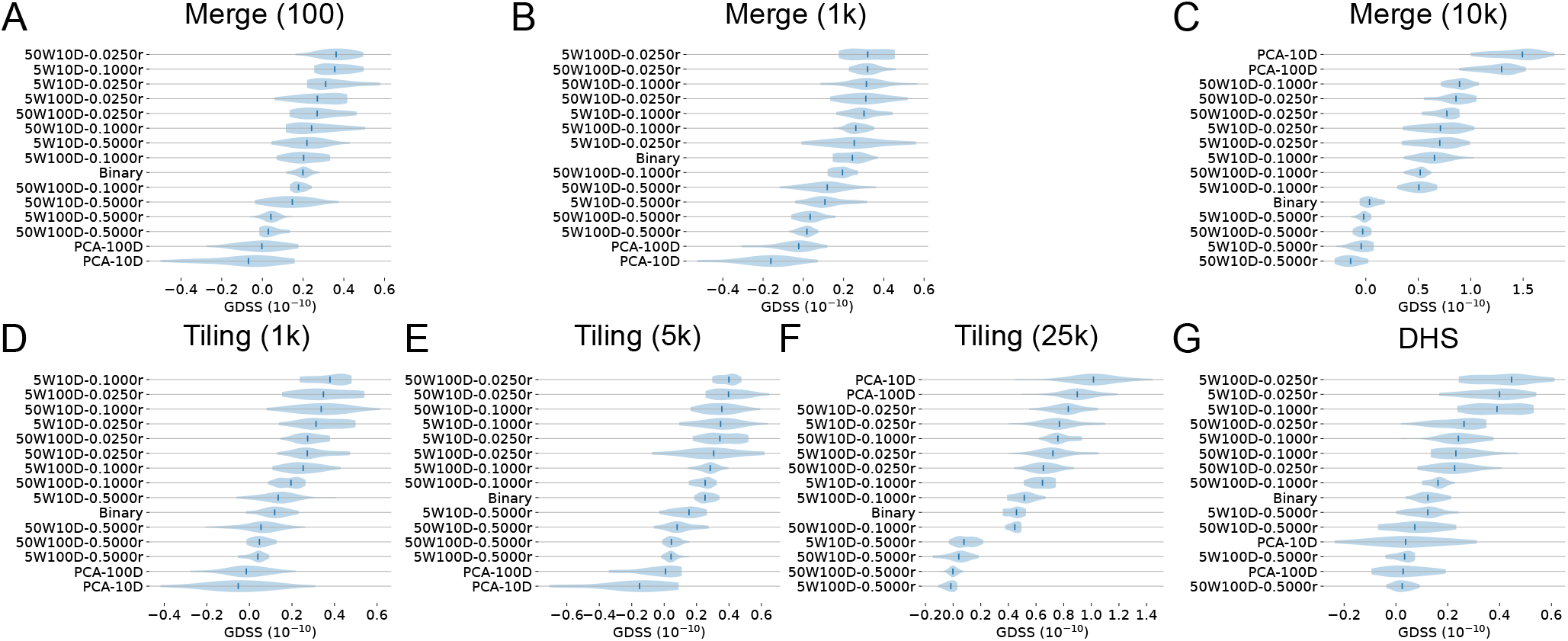
GDSSs of different sets of embeddings for regions in the seven universes. Each blue horizontal bar and the associated black bar indicate the average and standard deviation of the GDSSs calculated with 20 different random seeds for each set of region embeddings. W: context window size, D: embedding dimension, r: initial learning rate.

**Figure S4.**
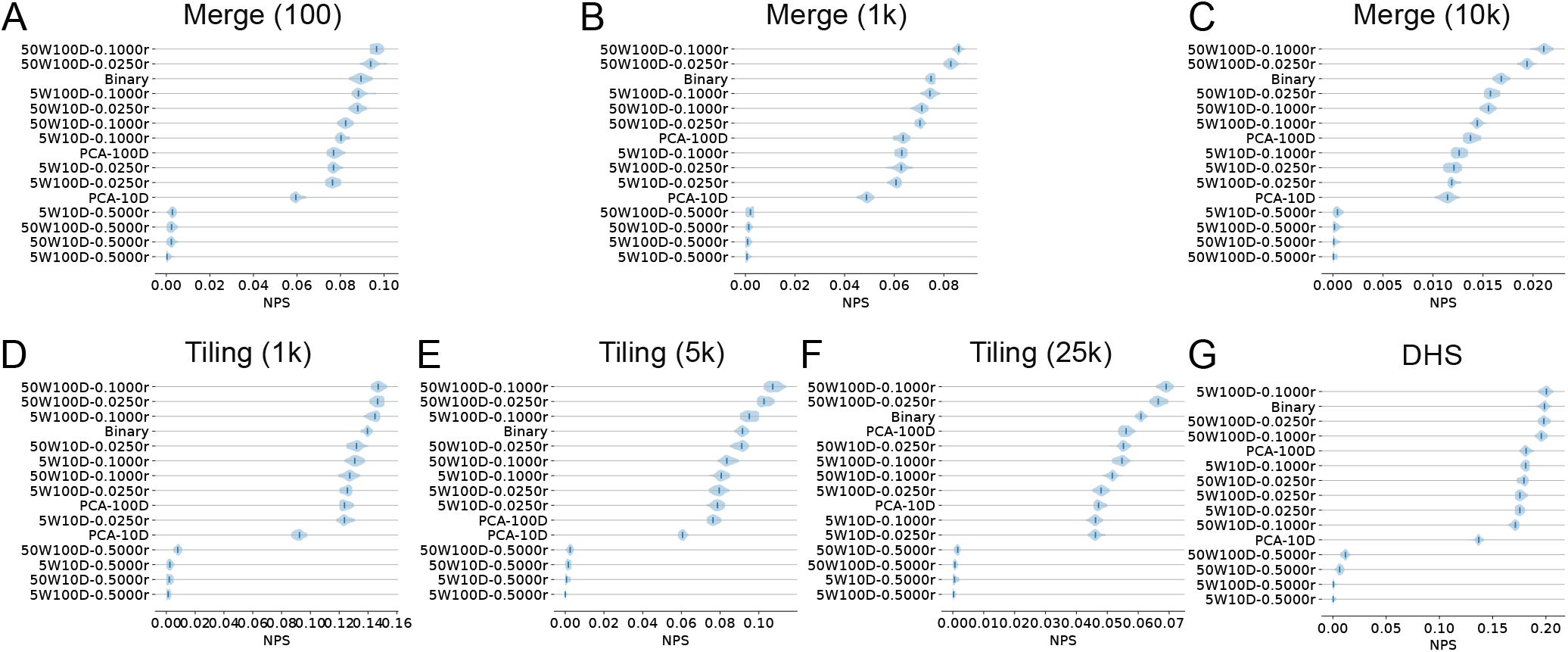
NPSs of different sets of embeddings for regions in the seven universes. Each blue horizontal bar and the associated black bar indicate the average and standard deviation of the NPSs calculated with 20 different random seeds for each set of region embeddings. W: context window size, D: embedding dimension, r: initial learning rate.

**Figure S5.**
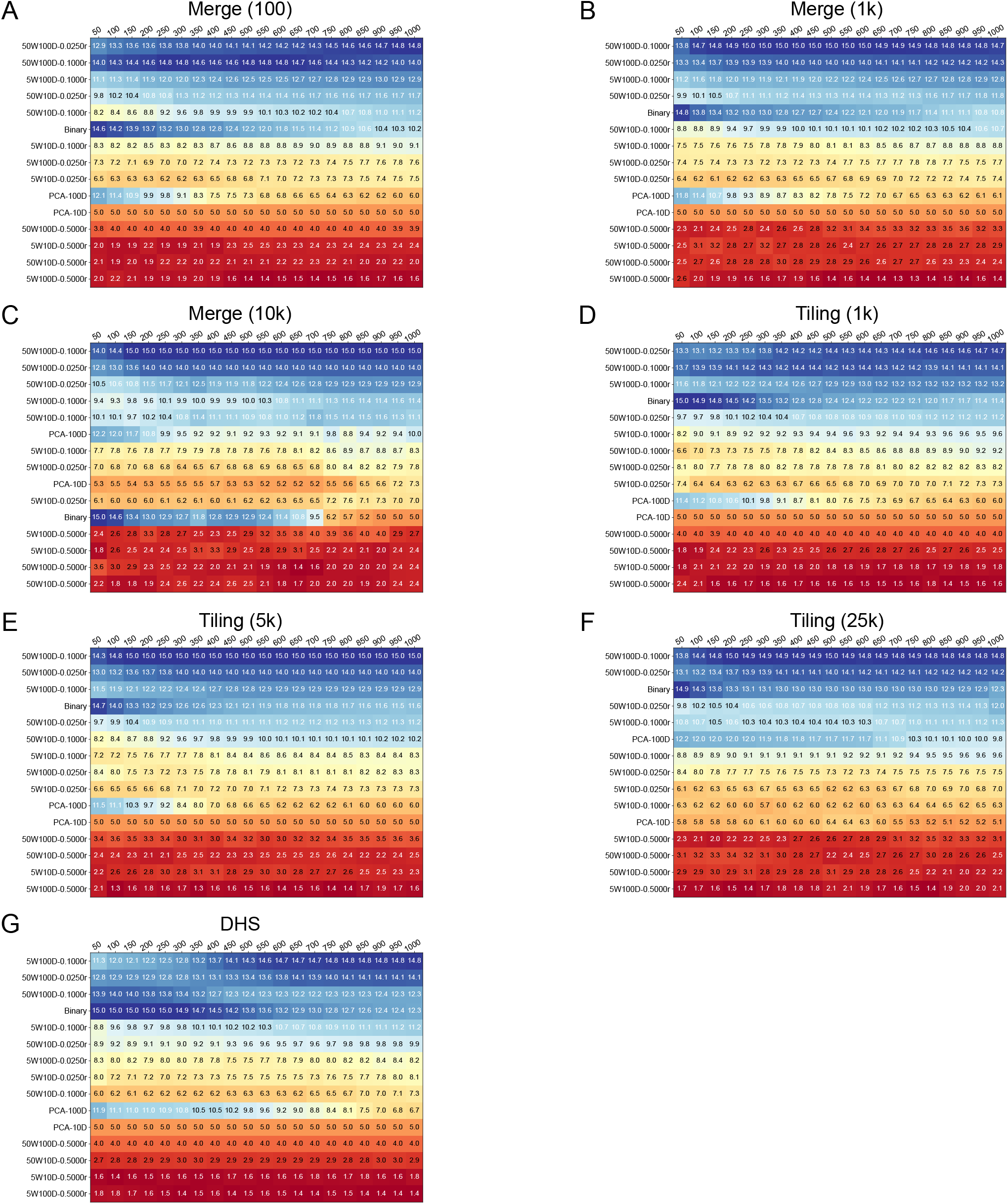
Average ranks of each set of region embeddings under different Us over 20 runs of the calculation of NPS. A large rank indicates a high NPS. W: context window size, D: embedding dimension, r: initial learning rate.

**Figure S6.**
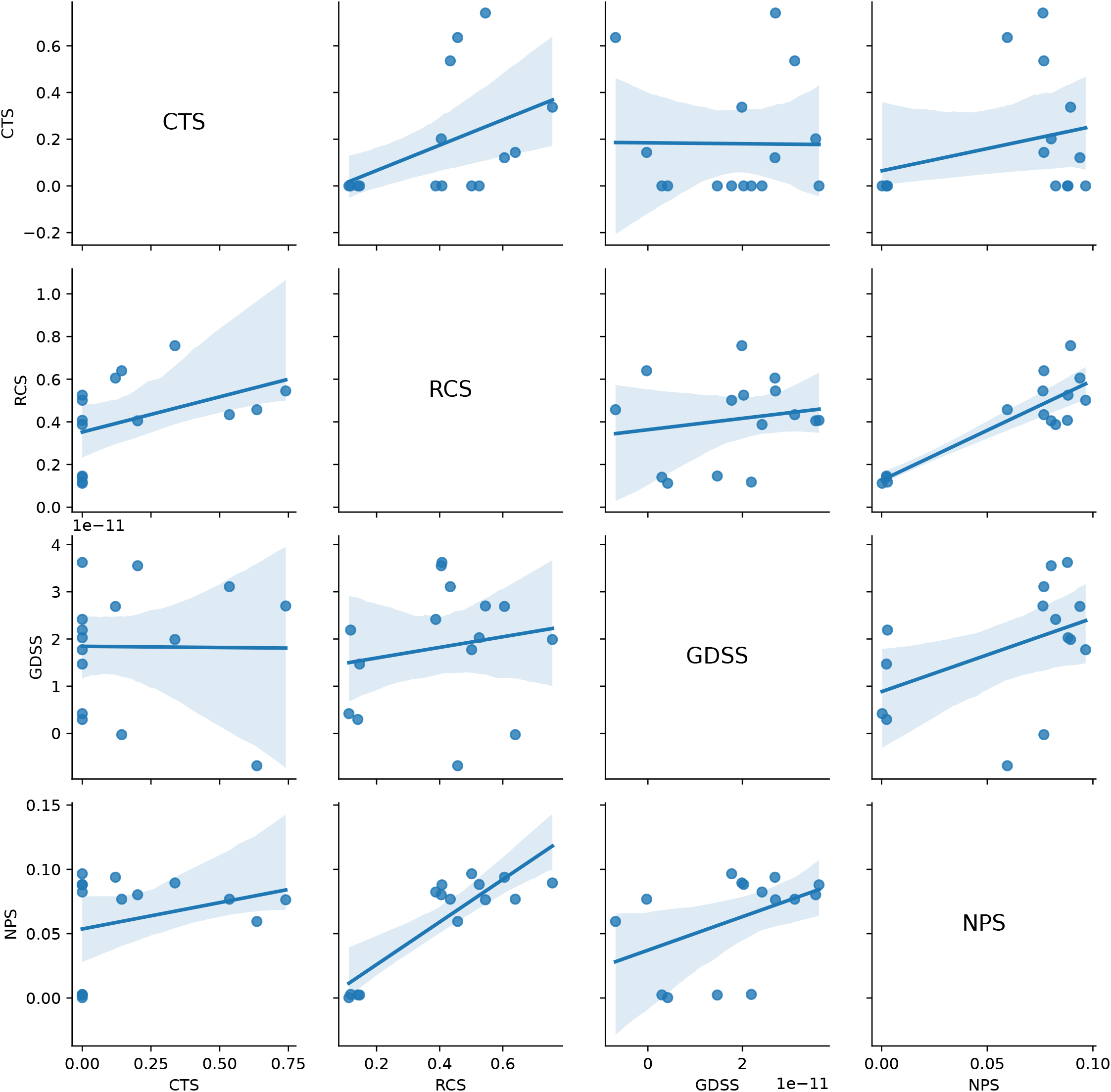
Pairwise correlations between the CTS, RCS, GDSS, and NPS for the Merge (100) universe.

